# Epigenetic regulation of p63 blocks squamous-to-neuroendocrine transdifferentiation in esophageal development and malignancy

**DOI:** 10.1101/2023.09.09.556982

**Authors:** Yongchun Zhang, Dimitris Karagiannis, Helu Liu, Mi Lin, Yinshan Fang, Ming Jiang, Xiao Chen, Supriya Suresh, Haidi Huang, Junjun She, Feiyu Shi, Patrick Yang, Wael El-Rifai, Alexander Zaika, Anthony E. Oro, Anil K. Rustgi, Timothy C. Wang, Chao Lu, Jianwen Que

## Abstract

While cell fate determination and maintenance are important in establishing and preserving tissue identity and function during development, aberrant cell fate transition leads to cancer cell heterogeneity and resistance to treatment. Here, we report an unexpected role for the transcription factor p63 (Trp63/TP63) in the fate choice of squamous versus neuroendocrine lineage in esophageal development and malignancy. Deletion of *p63* results in extensive neuroendocrine differentiation in the developing mouse esophagus and esophageal progenitors derived from human embryonic stem cells. In human esophageal neuroendocrine carcinoma (eNEC) cells, p63 is transcriptionally silenced by EZH2-mediated H3K27 trimethylation (H3K27me3). Upregulation of the major p63 isoform ΔNp63α, through either ectopic expression or EZH2 inhibition, promotes squamous transdifferentiation of eNEC cells. Together these findings uncover p63 as a rheostat in coordinating the transition between squamous and neuroendocrine cell fates during esophageal development and tumor progression.

## Introduction

Cell fate determination is critical for tissue development and homeostasis. Aberrant differentiation can lead to ectopic emergence of cells that are incompatible with organ function. For example, the aberrant presence of acid-secreting cells in the esophagus contributes to pathological lesions known as inlet patches.(*1*) Additionally, abnormal cell type conversions are found in premalignant lesions such as Barrett’s metaplasia where the stratified squamous epithelium is replaced by simple columnar cells in the distal portion of the esophagus (*2–4*). Chemotherapy and targeted therapy treatments have also been shown to induce a switch in cancer histology, such as androgen deprivation-induced prostate adenocarcinoma conversion into squamous cell carcinoma (*5, 6*). Moreover, neuroendocrine transdifferentiation was identified as a potential cause of drug resistance in several cancer types, including prostate adenocarcinoma and rectal adenocarcinoma (*7, 8*). Chemoradiotherapy can also cause esophageal cancer to switch from squamous cell carcinoma (SCC) to neuroendocrine carcinoma (*9*). Despite an increasing appreciation of the prevalence and significance of such tumor lineage plasticity, the underlying molecular mechanisms remain largely unknown.

The adult esophagus is lined with a stratified squamous epithelium, which starts as simple columnar cells in the early foregut (*10, 11*). Genetic studies have shown that the Bone Morphogenetic Protein (BMP) pathway plays a critical role in this transformation, involving the specification of basal progenitor cells into squamous epithelium (*12*). Downstream of BMP signaling, p63 (Trp63 in mouse, TP63 in human) is a key master transcription factor specifically enriched in basal progenitor cells (*13*). Deletion of *p63* blocks the formation of stratified squamous epithelium in the esophagus and skin (*14*). In skin keratinocytes, p63 has been shown to modulate the epigenomic landscape through recruitment of various chromatin regulators (*15–17*). Although abundant ciliated cells and mucous producing cells have been reported in the esophagus of *p63* null mutants (*13, 18*), how p63 is involved in cell fate determination in the esophageal progenitor cells remains incompletely understood.

Neuroendocrine cells play important roles in chemoreception, mechanotransduction and immune cell responses in the stomach, intestine and lung (*19–21*). Although Merkel cells, one type of neuroendocrine cell, have been detected in the human esophagus (*22*), their function remains unknown. In the mouse esophagus, existence of neuroendocrine cells has not been reported. Notably, neuroendocrine carcinoma also occurs in the esophagus, accounting for 0.4–2.8% of human esophageal cancers (*23*). Esophageal neuroendocrine carcinoma (eNEC) is highly aggressive and composed of cancer cells characterized by high expression levels of neuroendocrine cell markers such as SYP and CHGA, similar to small-cell lung cancer (SCLC) (*24, 25*). It is noteworthy that eNECs are negative for the expression of p63 (*26*), yet genome sequencing reveals no mutation in the gene encoding p63 (*27*), suggesting an epigenetic mechanism underlying p63 silencing during tumor progression.

Histone posttranslational modifications such as methylation and acetylation modulate the structure of chromatin and impact gene transcription. The trimethylation of lysine 27 on histone H3 (H3K27me3) is catalyzed by the methyltransferase EZH2, a core subunit of Polycomb Repressive Complex 2 (PRC2) (*28*). H3K27me3 generally represses transcription of genes that are essential for lineage specification (*29*). Hence, EZH2 is important for tissue development, homeostasis and tumorigenesis (*28*). By contrast, histone acetylation generally activates gene transcription (*30*). Histone acetylation is regulated by both histone acetyltransferase (e.g., p300/CBP) and histone deacetylases (HDACs) (*31*). In addition, the Bromodomain and Extraterminal (BET) protein family including BRD2, BRD3, BRD4, and BRDT recognize and bind to acetylated histone lysine residues to recruit RNA polymerase II to promote gene transcription (*32*). Accordingly, histone acetylation markers such as H3K27 acetylation (H3K27ac) demarcate active *cis*-regulatory elements critical for tissue development (*33*).

Here, we report that epigenetic regulation of p63 is critical for esophageal development and tumor lineage plasticity. p63 acts as a conserved guardian against neuroendocrine cell fate, and *p63* deletion leads to abundant neuroendocrine cell differentiation in the developing esophagus. Conversely, p63 overexpression confers squamous cell differentiation in eNEC cells. We further show that p63 expression is epigenetically silenced by H3K27me3 in eNEC cells. Treatment with EZH2 inhibitors reduces H3K27me3, leading to the expression of p63 and squamous identity genes in eNEC organoids.

## Results

### p63 represses neuroendocrine cell differentiation in the developing mouse esophagus

We and others have shown that deletion of *p63* causes the emergence of mucociliated cells in the esophagus (*3, 13, 18*). To comprehensively identify the cell types present in the *p63*-null developing esophagus, we compared gene expression of the esophageal epithelium between *p63 KO* mutants and the littermate *wild-type (WT)* controls at E12.5 (**Fig. 1A**). Differential expression analysis revealed that 954 and 703 genes were significantly upregulated and downregulated in *p63 KO* mutants, respectively (**Fig. 1B**). Gene ontology analysis and gene set enrichment analysis (GSEA) revealed a significant impact of p63 loss on cell fate pathways and related biological processes (**Fig. 1, C to E, and fig. S1A**). Specifically, we observed downregulation of genes associated with basal and squamous cell identity and extracellular matrix organization (**Fig. 1, C and D, and fig. S1A**). The basal cell signature genes *Krt5* and *Krt15*, and adhesion molecules including *Itga3, Itga6, Itgb4* and *Lamb3* were reduced in the mutant esophageal epithelium (**Fig. 1F**). As expected, the expression levels of the ciliated genes *Tuba1a* and *Tubb4a*, and the columnar cell keratins *Krt7, Krt8* and *Krt18* were significantly increased upon *p63* deletion (**Fig. 1F**). By contrast, we found that genes upregulated in mutants were enriched for those regulating neuroendocrine cell identity and neurotransmitter processes (**Fig. 1, C and E**). We observed upregulation of neuroendocrine cell signature genes including *Ascl1*, *Insm1*, *Cgrp*, *Syp*, *Chga*, *Chgb*, *Edn1*, *Eno2*, *Cck*, *Uch1*, *Gfi1,* and *Ncam1* in mutants (**Fig. 1F**). We further used immunofluorescence (IF) staining to confirm the ectopic presence of INSM1^+^ neuroendocrine progenitor cells in the mutant but not wildtype esophagus (**Fig. 1G**). We also observed abundant mature neuroendocrine cells expressing CGRP and SYP in the mutant esophagus at E18.5 (**Fig. 1, H and I, and fig. S1B**). These data confirm that *p63* deletion leads to the ectopic presence of neuroendocrine cells, suggesting that p63 is required for protecting against aberrant neuroendocrine differentiation during esophageal development.

**Fig. 1.**
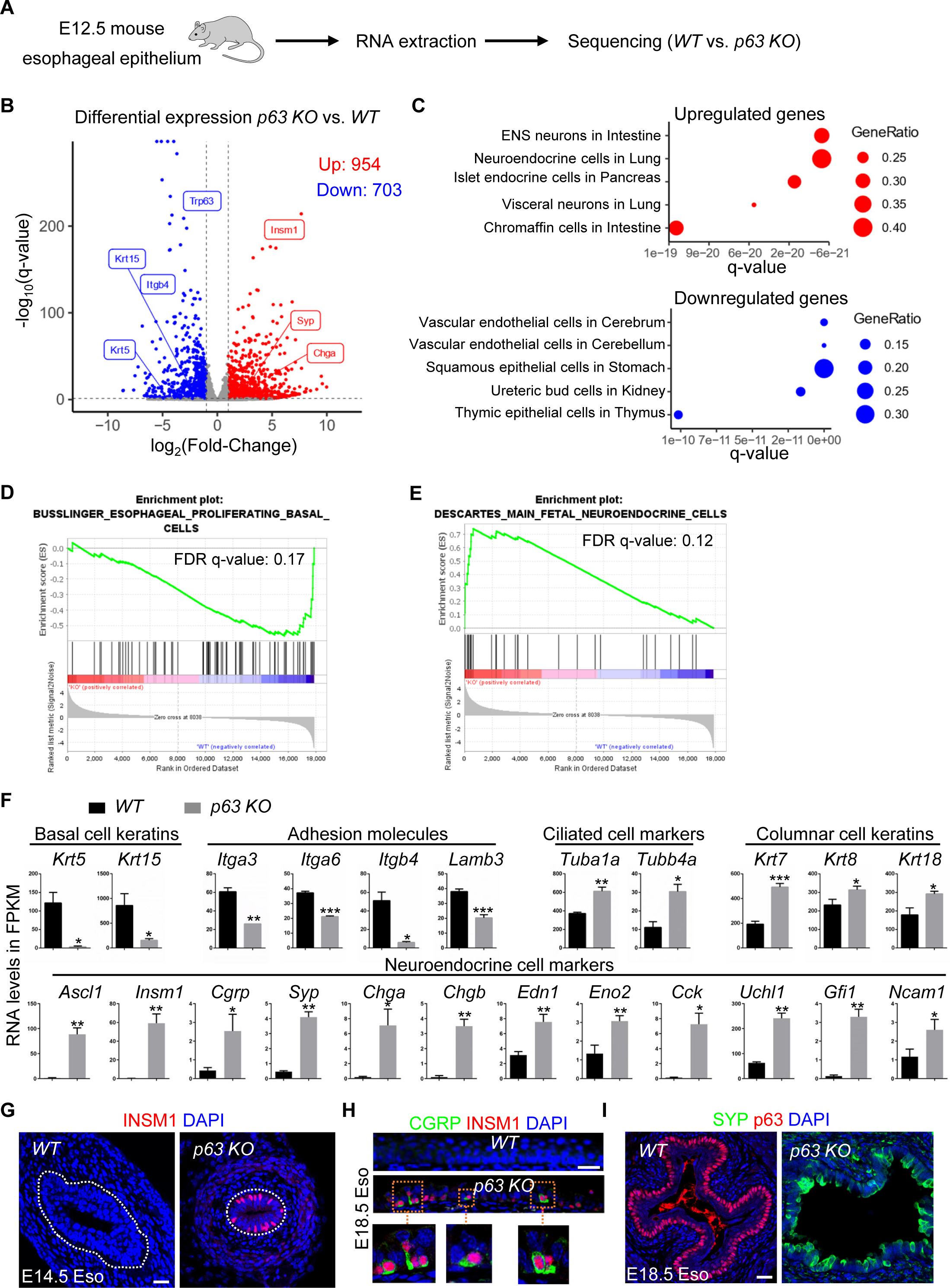
Loss of p63 leads to neuroendocrine cell differentiation in the developing mouse esophagus. **(A)** Schematics showing the epithelial isolation, RNA extraction and sequencing of the esophagus from *WT* and *p63 KO* mice. **(B)** Volcano plot of differentially expressed genes between *WT* and *p63 KO* mouse esophageal epithelium. Noted are genes involved in squamous and neuroendocrine cell identity. **(C)** Cell types and tissues gene ontology of genes upregulated (red) and downregulated (blue) in *p63 KO* mouse esophageal epithelium annotated by Descartes Cell Types and Tissues 2021. **(D-E)** Gene set enrichment analysis (GSEA) indicating downregulation of the esophageal basal cell signature and upregulation of the neuroendocrine cell signature. **(F)** *p63* deletion reduces the transcript levels of the basal cell keratins *Krt5* and *Krt15*, the adhesion molecules *Itga3*, *Itga6*, *Itgb4*, and *Lamb3,* but increases the levels of the ciliated cell genes *Tuba1a* and *Tubb4a*, and neuroendocrine cell signature genes including *Ascl1*, *Insm1*, *Cgrp*, *Syp*, *Chga*, *Chgb*, *Edn1*, *Eno2*, *Cck*, *Uch1*, *Gfi1,* and *Ncam1.* The esophagi examined are at E12.5. *p < 0.05, **p < 0.01, ***p < 0.001. **(G)** The neuroendocrine cell marker INSM1 is ectopically expressed in the epithelium of E14.5 *p63 KO* esophagus. **(H-I)** The neuroendocrine markers INSM1, CGRP and SYP are expressed in the epithelium of *p63 KO* esophagus. Abbreviation: Eso, esophagus. Scale bars: 20 µm.

### p63 plays a conserved role in repressing neuroendocrine cell fate in human ESC-derived esophageal progenitor cells

To address whether p63 plays a similar role in inhibiting neuroendocrine cell fate in human esophageal progenitor cells (EPCs), we deleted *p63* in the H9 human embryonic stem cell (hESC) line. We then differentiated H9 into EPCs in a 2D system using the protocol we previously described (*34, 35*). Remarkably, we detected increased expression of the neuroendocrine genes *ASCL1, INSM1, SYP, CGRP, CHGA*, and *SST* at day 24 of differentiation (**Fig. 2A**), and ASCL1^+^ neuroendocrine cells were present among EPCs following *p63* deletion (**Fig. 2B**). In addition, we embedded endodermal spheroids in Matrigel and induced them to form 3D esophageal organoids (**Fig. 2C**). We collected the organoids after 7 weeks in culture and determined the expression of genes marking basal cells versus neuroendocrine cells. The expression of p63, KRT5 and KRT13 was absent in the p63 *KO* organoids in comparison to their high levels of expression in the *WT* organoids (**Fig. 2, D to E**). Importantly, we observed the ectopic presence of INSM1^+^SYP^+^ neuroendocrine cells in organoids formed by *p63*-null hESC-derived EPCs (**Fig. 2F**). By contrast, INSM1^+^SYP^+^ cells were not detected in the organoids formed by control hESC-derived EPCs (**Fig. 2F**). Taken together, these findings suggest a conserved role for p63 in repressing neuroendocrine cell differentiation during the development of the human esophagus.

**Fig. 2.**
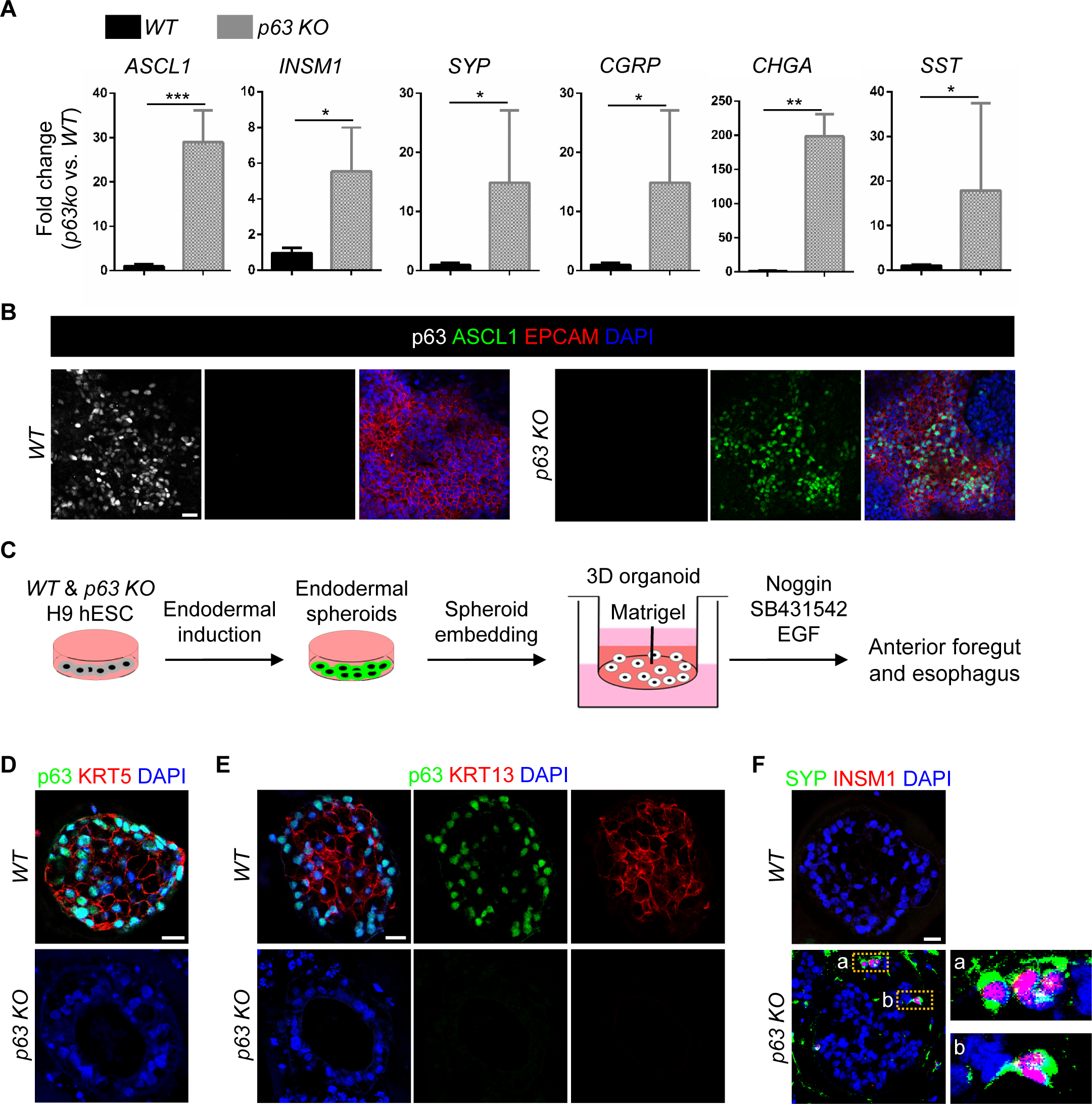
p63 deletion leads to neuroendocrine differentiation of esophageal progenitor cells derived from human embryonic stem cells (hESCs) **(A-B)** p63 deletion leads to ectopic neuroendocrine cell differentiation, shown by the increased transcript levels of *ASCL1*, *INSM1*, *SYP*, *CGRP*, *CHGA*, and *SST* (A) and ectopic expression of ASCL1 protein (B). Black bar, *WT*; Grey bar, *p63 KO.* (C) Schematics showing the differentiation of H9 hESCs into esophageal progenitor cells and the formation of esophageal organoids. Endodermal organoids derived from H9 hESCs cells were embedded in Matrigel and cultured in the presence of Noggin, SB431542, and EGF for 7 weeks to form esophageal organoids. (D) *p63* deletion leads to the loss of KRT5 expression in hESC-derived esophageal cells. (E) *p63* deletion perturbs the formation of stratified esophageal organoids. Note that the WT organoids express high levels of p63 and KRT13 in the periphery and center, respectively. The expression pattern is lost in *p63 KO* organoids. **(F)** p63 deletion leads to the ectopic expression of INSM1 and SYP (yellow boxes) in hESC-derived esophageal organoids. *KO*, *knockout*. Scale bars: 20 µm.

### p63 expression is repressed by EZH2-mediated H3K27 methylation in eNECs

Because *p63* deletion led to the ectopic presence of neuroendocrine cells in the esophagus, we hypothesized that p63 inactivation is required to maintain esophageal neuroendocrine cell identity. While neuroendocrine cells have yet to be identified in the mouse esophagus, Merkel cells have been detected in the human esophagus. Notably, p63 expression is lost in human esophageal neuroendocrine carcinomas (eNEC) (*24, 26*). We confirmed that p63 was not detected in INSM1^+^ neuroendocrine cancer cells in human eNEC specimens, although neighboring basal cells expressed high levels of p63 (**Fig. 3A**). Mutational profiling indicated that *p63* is not genetically mutated in a large cohort of eNECs (*27*). We therefore asked whether p63 is silenced through epigenetic mechanisms.

**Fig. 3.**
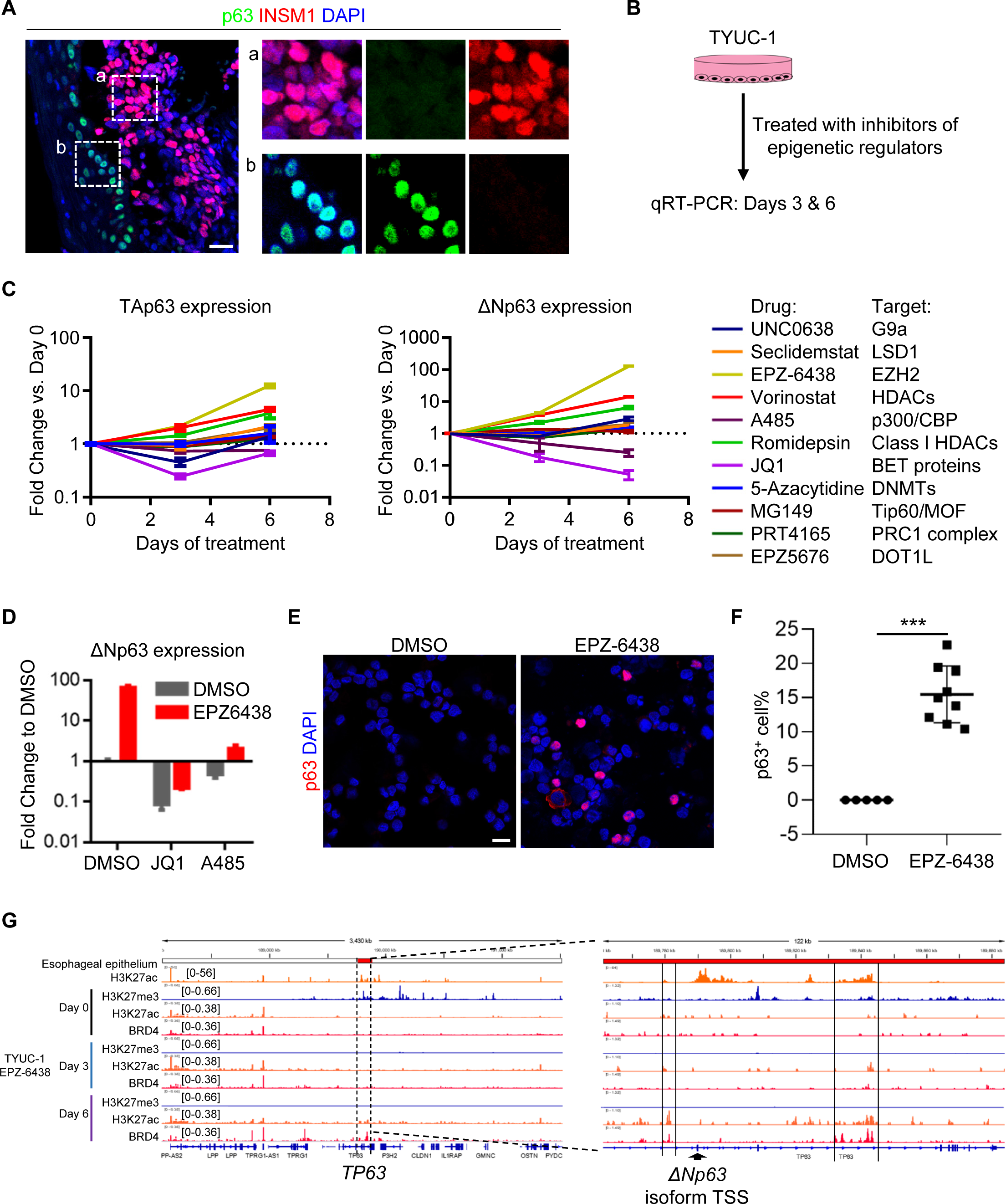
ΔNp63α expression is regulated by methylation and acetylation of histone H3 lysine 27. **(A)** p63 and INSM1 staining in human esophageal small cell carcinoma specimens. Scale bars: 20 µm. **(B)** Schematics showing that TYUC-1 esophageal neuroendocrine cancer cells were treated with inhibitors of epigenetic regulators and gene expression was analyzed by qRT-PCR. **(C)** The transcript levels of TAp63 and ΔNp63 in TYUC-1 esophageal neuroendocrine cancer cells treated with various epigenetic inhibitors. Note that the EZH2 inhibitor EPZ-6438 increased the expression of TAp63 and ΔNp63 to the highest levels compared to other inhibitors. **(D)** Transcript levels of ΔNp63 in TYUC-1 cells treated with EPZ-6438 alone or in combination with JQ1 or A485. **(E-F)** Immunofluorescence staining and quantification of p63 cells in EPZ-6438 treated TYUC-1 cells. Note that p63 is expressed in the EPZ-6438-treated group but not the DMSO-treated group. Scale bars: 20 µm. **(G)** Genome browser view at the *TP63* locus showing esophagus epithelium H3K27ac ChIP-seq (ENCODE SRX3205374) and TYUC-1 cell H3K27me3, H3K27ac, and BRD4 CUT&Tag upon treatment with EPZ-6438 for 3 and 6 days. Noted are putative enhancer elements near the ΔNp63 isoform TSS where H3K27 methylation is lost, and acetylation is restored.

The *TP63* gene encoding human p63 harbors two alternative promoters, resulting in TAp63 isoforms with the N-terminal transactivation (TA) domain, or ΔNp63 isoforms lacking the TA domain. We first tested whether TAp63 and ΔNp63 expression are modulated epigenetically by screening a panel of chromatin regulator inhibitors in a human eNEC cell line (TYUC-1) (**Fig. 3B and fig. S2**). We found that inhibition of EZH2, the histone methyltransferase that catalyzes H3K27me3 deposition, with the FDA-approved small molecule inhibitor EPZ-6438 induced the highest expression levels of ΔNp63 (**Fig. 3C**), indicating that H3K27me3 represses p63 expression in eNECs. In addition, inhibition of histone deacetylases (HDACs) with either Vorinostat or Romidepsin increased ΔNp63 expression (**Fig. 3C**). EZH2 and HDAC inhibitors also modestly upregulated the expression of TAp63 (**Fig. 3C**). Conversely, inhibition of histone acetyltransferase p300/CBP and BET proteins, “readers” of histone lysine acetylation, reduced TAp63 and ΔNp63 expression (**Fig. 3C**). These results suggest that the post-translational modification state of H3K27 (acetylation vs. methylation) has a direct effect on p63 expression. Consistently, the transcriptional activating effect of EZH2 inhibition (EPZ-6438) on ΔNp63 was abolished by co-treatments with either BET protein (JQ1) or CBP/p300 (A485) inhibitors (**Fig. 3D**), suggesting that histone acetylation and BET protein binding are required for ΔNp63α expression following H3K27me3 depletion. We also found that EPZ-6438 treatment increased the protein abundance of p63 in TYUC-1 cells (**Fig. 3, E and F**).

To assess the chromatin landscape following EZH2 inhibition, we performed Cleavage Under Targets and Tagmentation (CUT&Tag) of H3K27me3, H3K27ac and BRD4 upon EPZ-6438 treatment (**Fig. 3G**). As expected, H3K27me3 was enriched in the *TP63* locus and was lost upon EPZ-6438 treatment in TYUC-1 cells (**Fig. 3G**). Moreover, EPZ-6438-treated cells gained two peaks of H3K27ac and BRD4 binding at putative intronic enhancers close to the transcription start site (TSS) of ΔNp63 isoforms (**Fig. 3G, right panel**). Notably, re-analysis of ChIP-seq data of esophageal epithelium from ENCODE indicated the presence of H3K27ac peaks at these sites, indicating that they represent physiologically relevant squamous-specific *cis*-regulatory elements associated with p63 expression (**Fig. 3G**). Taken together these results suggest that epigenetic modifications mediated by EZH2 play a critical role in silencing p63 in eNECs.

### ΔNp63α is the major isoform driving squamous transdifferentiation program in eNEC cells

To investigate whether restoration of p63 expression impacts neuroendocrine cell identity, we first used a doxycy higher transcription of genes cline-inducible system to overexpress p63 in TYUC-1 cells (**Fig. 4A**). We chose to express two major p63 isoforms, TAp63α and ΔNp63α (*36*), both present in the esophagus, to determine their effects on altering the cell fate of eNEC. RNA-sequencing indicated that ectopic ΔNp63α expression induced higher transcription of genes associated with squamous identity than TAp63α (**Fig. 4B**). To exclude the possibility that the differential induction of squamous identity genes is due to the differences in p63 isoform expression levels, we measured KRT5 expression at a concentration of doxycycline where TAp63α and ΔNp63α were induced at comparable levels. We confirmed that ΔNp63α was more potent in activating KRT5 expression (**Fig. 4C**), which is consistent with the dominant role of ΔNp63 over TAp63 in the regulation of keratin gene expression during keratinocyte differentiation (*37*).

**Fig. 4.**
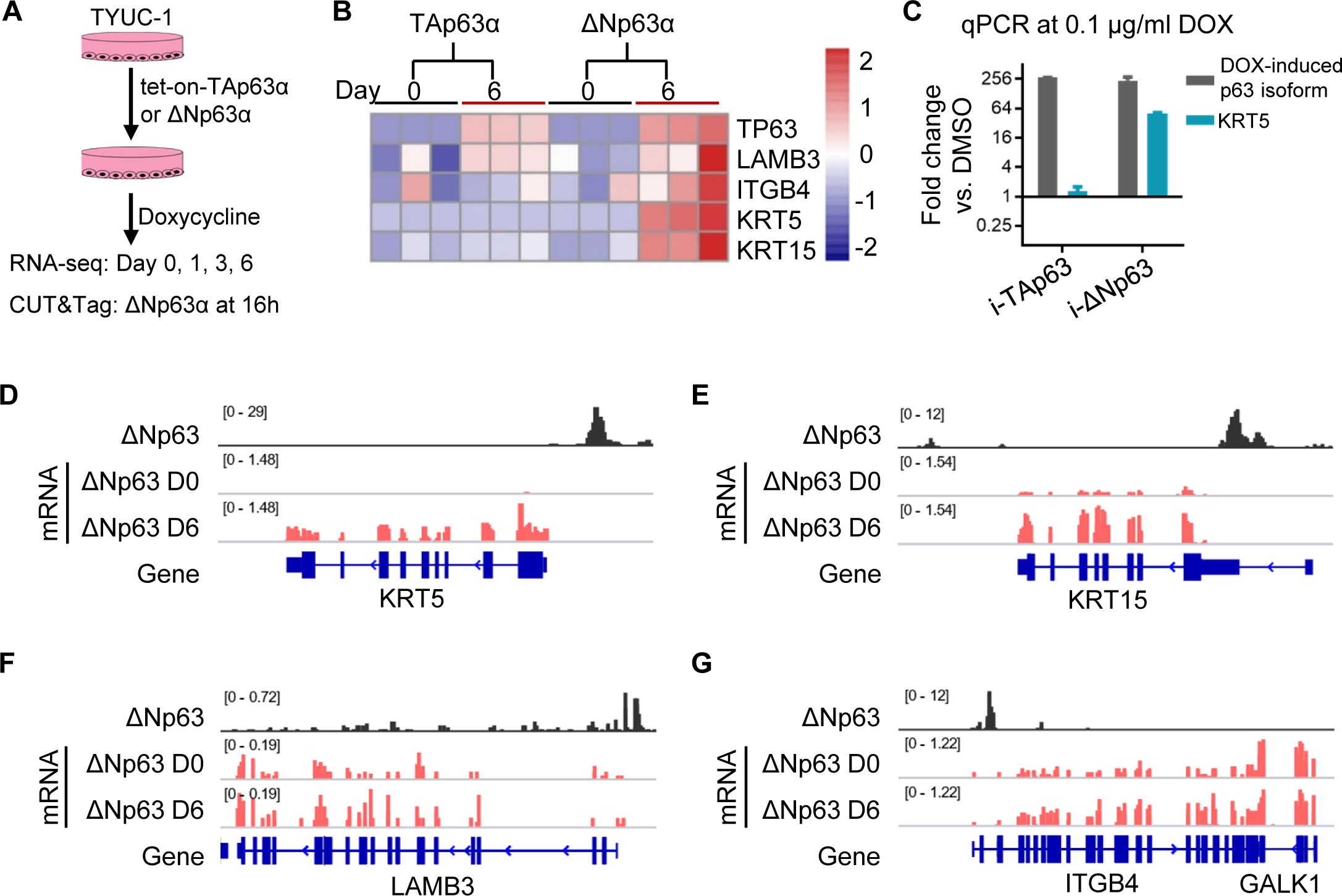
When overexpressed ΔNp63 is the major isoform that controls squamous identity genes in esophageal neuroendocrine cancer cells. **(A)** Schematics showing overexpressing ΔNp63α and TAp63α in TYUC-1 cells. Gene expression was analyzed by RNA-seq, and ΔNp63α binding sites on chromatin were profiled by CUT&Tag assays. **(B)** A heatmap of the transcript levels of squamous cell markers. Note that *KRT5* and *KRT15* genes were only induced by ΔNp63. **(C)** RT-qPCR of inducible *p63* isoforms and *KRT5* at a concentration of doxycycline where TAp63 and ΔNp63α were induced at comparable levels. DOX, doxycycline. **(D-G)** Genome browser view of p63 CUT&Tag and RNA-seq upon induction of ΔNp63 expression. Note that ΔNp63 directly binds to *LAMB3*, *ITGB4*, *KRT5,* and *KRT15*.

To determine if ectopically expressed ΔNp63α directly activates squamous genes in eNEC cells, we assessed ΔNp63α genomic distribution using CUT&Tag in TYUC-1 cells where ΔNp63α expression was induced for 16 hours. Motif analysis of ΔNp63α peaks confirmed enrichment for the p63 binding motif as expected (**fig. S3A**). Peak annotation indicated that the majority of ΔNp63α peaks were found within gene promoters and introns (**fig. S3B**), consistent with the notion that p63 regulates gene expression through binding at *cis*-regulatory elements (*38*). Importantly, ΔNp63α peaks were identified close to the promoters of the squamous identity genes including *KRT5*, *KRT15*, *LAMB3,* and *ITGB4*, suggesting a direct regulation of their expression (**Fig. 4, D to G**). These results indicate that ΔNp63α but not TAp63α is the major isoform that upregulates squamous genes upon overexpression in eNEC cells through direct binding at their promoters.

We further characterized the transcriptomic changes in TYUC-1 cells following ΔNp63α overexpression for 6 days (**Fig. 5A**). Differential gene expression analysis indicated that more genes were upregulated than downregulated (207 vs. 41), suggesting a positive role for ΔNp63α in gene expression (**Fig. 5A**). Moreover, we observed an enrichment of squamous genes in the upregulated gene list (**Fig. 5B and fig. S4A**), as well as an increasing predominance of squamous gene expression signature over time (**Fig. 5C**). Notably, among 5877 genes that were associated with at least one ΔNp63α peak, less than 2% (90) were upregulated upon ΔNp63α overexpression (**Fig. 5D**), suggesting that ΔNp63α binding alone is often not sufficient to increase the expression of its target genes. This result is in agreement with previous work showing that additional chromatin elements such as the presence of histone H3 acetylation are required for efficient induction of gene expression by p63 (*38*). Furthermore, we overexpressed ΔNp63α in TYUC-1 cells cultured as 3D tumor organoids. ΔNp63α overexpression resulted in the ectopic protein expression of KRT5 and KRT15 (**Fig. 5, E and F, and fig. S4B**), and the differentiated squamous cell markers KRT4 and KRT13 (**Fig. 5, G and H**). Together, these results suggest that ectopic ΔNp63α expression confers squamous cell identity to eNEC cells.

**Fig. 5.**
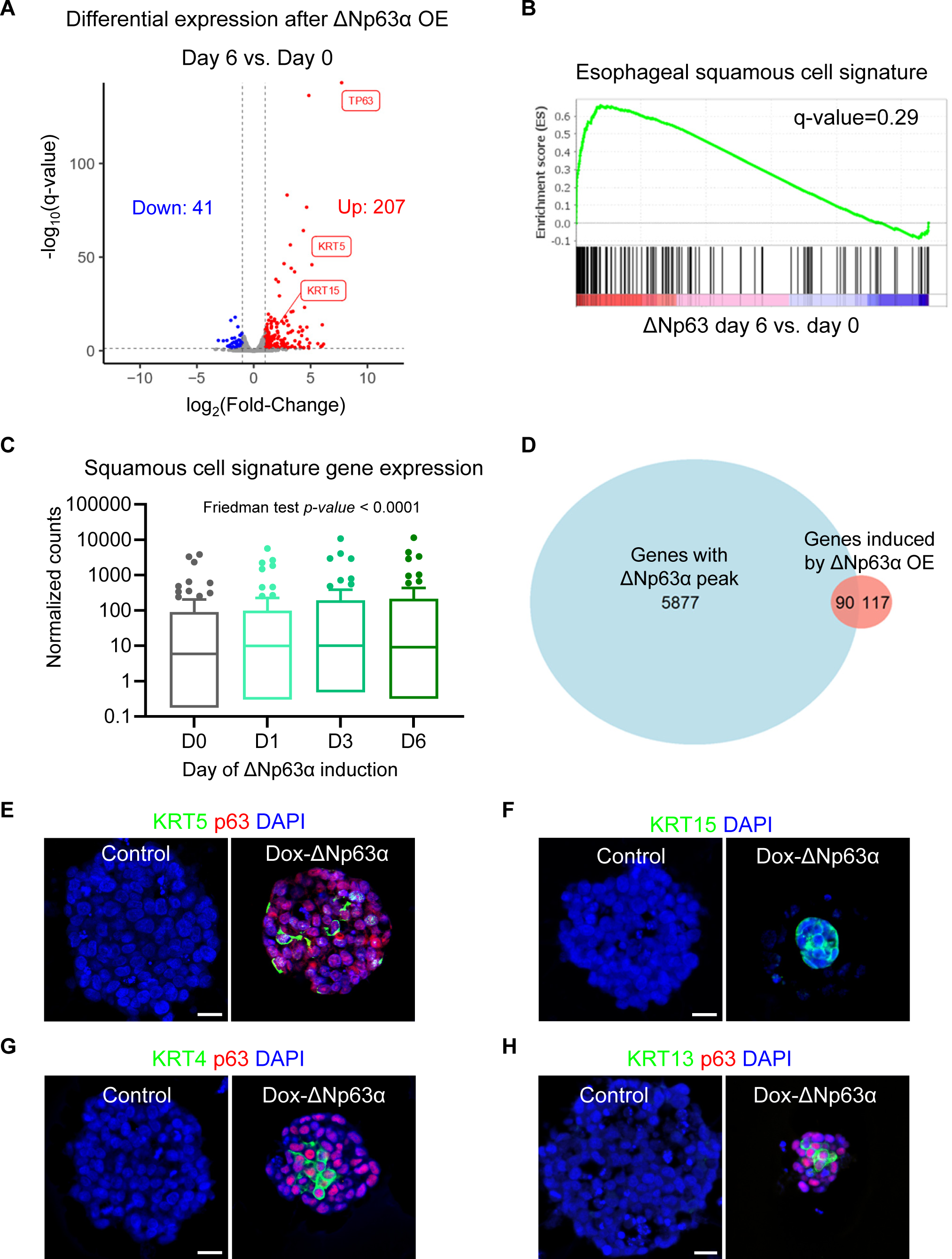
ΔNp63 reprograms esophageal neuroendocrine cancer cells into squamous cells. **(A)** Volcano plot of differentially expressed genes upon induction of ΔNp63α expression by doxycycline for 6 days. OE, overexpression. **(B)** GSEA analysis showing that the esophageal squamous cell gene signature is positively correlated with ΔNp63 overexpression over time. **(C)** Expression levels of genes of the ‘Descartes Fetal Stomach Squamous Epithelial Cells’ gene set at 0, 1, 3, and 6 days of doxycycline. **(D)** Venn diagram showing the overlap between ΔNp63α-bound genes and ΔNp63α-induced genes. OE, overexpression. **(E-H)** Immunofluorescence staining of p63, KRT5, KRT15, KRT4, and KRT13 in the TYUC-1 tumor organoids with doxycycline-induced overexpression of ΔNp63α (Dox-ΔNp63α) versus the untreated (Control). Scale bars: 20 µm.

### EZH2 inhibition promotes squamous differentiation of eNEC cells through de-repressing p63

We next determined if epigenetic reactivation of ΔNp63α through EZH2 inhibition similarly drives squamous transdifferentiation of eNEC cells. We performed RNA-sequencing and qRT-PCR to analyze gene expression of TYUC-1 cells following EPZ-6438 treatment for 6 days. EZH2 inhibition significantly induced the expression of *p63* and squamous cell genes including *KRT5*, *KRT14* and *SOX2* (**Fig. 6A and fig. S5A**). Furthermore, Gene Ontology Enrichment Analysis revealed that EZH2 inhibition promoted the expression of genes that are associated with squamous epithelial cells in tissues like the forestomach and lung (**Fig. 6B**). This chromatin and transcriptome remodeling was accompanied by increased apoptosis, as shown by the accumulation of cleaved caspase-3 (**fig. S5B**), as well as decreased cell proliferation (**fig. S5C**). Additionally, EPZ-6438 treatment decreased the frequencies (DMSO, 1.65 ± 0.06%; EPZ-6438, 0.84 ± 0.04%; p < 0.001) and sizes of eNEC organoids (DMSO, 192.35 ± 59.14 µm; EPZ-6438, 85.82 ± 19.67 µm; p < 0.001) (**fig. S5, D to F**). We also observed the expression of *p63, KRT5,* and *SOX2* in eNEC organoids treated with EPZ-6438 (**Fig. 6, C to E**). In contrast, the levels of H3K27me3 and the neuroendocrine marker *SYP* were decreased in p63^+^ cells (**Fig. 6, C and F, and fig. S5A**). Overall, p63 was ectopically expressed in 22.7% of organoids upon EZH2 inhibition (**Fig. 6G**), and 73.6% cells in these organoids expressed p63 (**Fig. 6H**). To rule out potential off-target effects of drug treatment, we used another EZH2 inhibitor, GSK126, and observed consistent reduction in the formation efficiency (DMSO, 1.71 ±0.06%; GSK126, 0.93 ±0.07%; p < 0.001) and size of eNEC organoids (DMSO, 189.76 ± 59.09 µm; EPZ-6438, 94.92 ± 19.67 µm; p < 0.001) (**fig. S5G**). Organoids treated with GSK126 also exhibited increased expression of p63 and KRT5 (**fig. S5, H and I**). Furthermore, we used two independent shRNAs to knockdown EZH2 in TYUC-1 organoids (**fig. S6, A and B**), and found that EZH2 knockdown consistently led to the ectopic expression of p63 and KRT5 (**fig. S6C**). To determine whether p63 is critical for the neuroendocrine-to-squamous transdifferentiation following EZH2 inhibition, we used siRNA to knockdown p63 and observed reduced upregulation of squamous genes upon EPZ-6438 treatment (**Fig. 6I**). Taken together, these results confirm that EZH2 epigenetically suppresses the expression of p63 in eNECs and that p63 governs the transition between squamous and neuroendocrine cell identities.

**Fig. 6.**
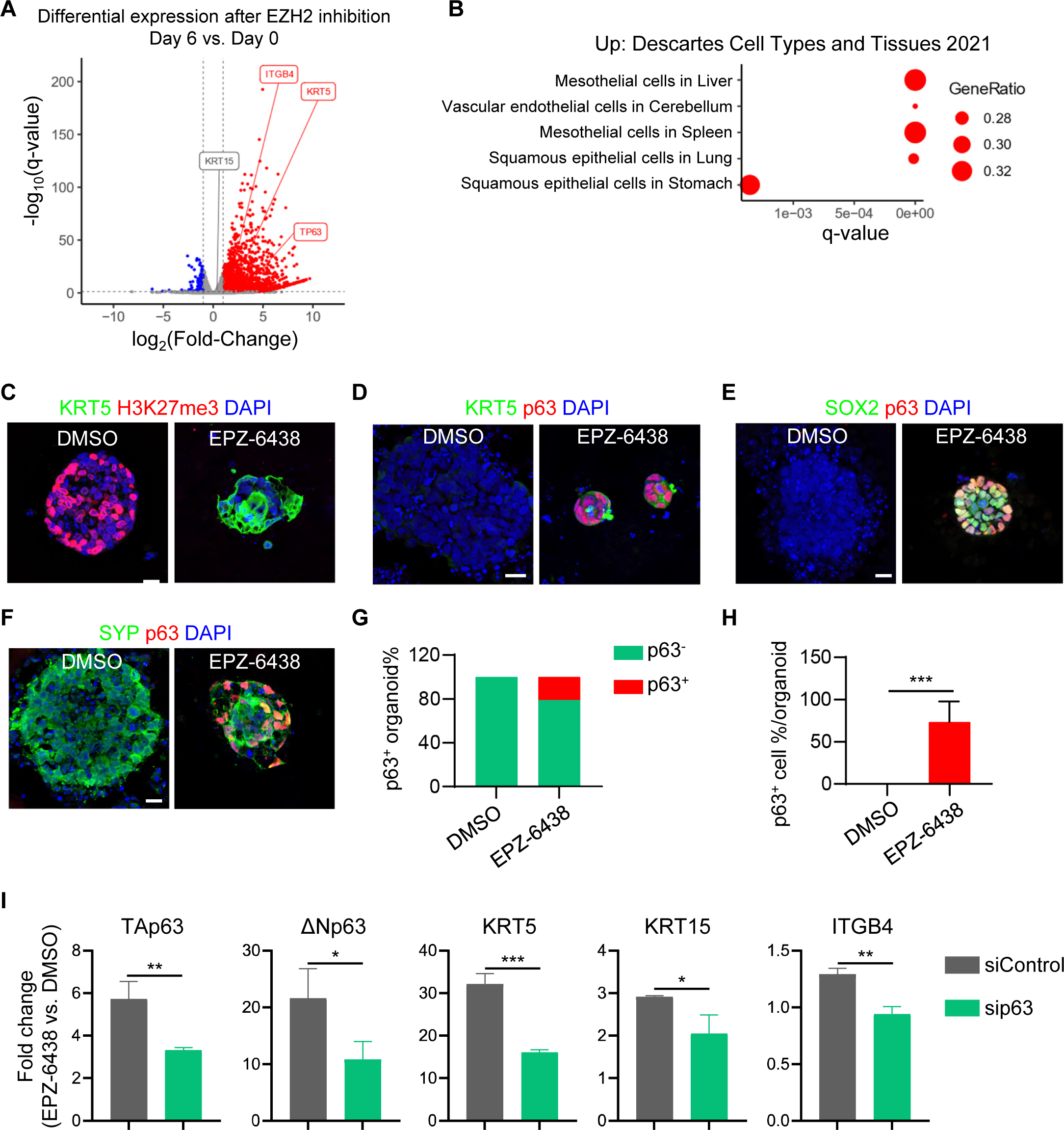
EZH2 inhibition promotes squamous differentiation of esophageal neuroendocrine cancer organoids through ΔNp63. **(A)** Volcano plot of differentially expressed genes upon treatment of TYUC-1 cells with DMSO or 2 μM EPZ-6438 for 6 days. **(B)** “Cell Types And Tissues” gene ontology enrichment analysis of upregulated genes. **(C-F)** Immunofluorescence staining of H3K27me3, basal cell markers p63, KRT5, and SOX2, and the neuroendocrine cell marker SYP in 3D TYUC-1 organoids. **(G-H)** Quantification of p63^+^ organoids (G) and p63^+^ cells (H) in organoids that contain p63^+^ cells. ***, p < 0.001. **(I)** Gene expression fold change (2 µM EPZ-6438 vs. DMSO) of TYUC-1 organoids transfected with p63 (sip63) or control (siControl) siRNA. *p < 0.05, **p < 0.01, ***p < 0.001. Scale bars: 20 µm.

## Discussion

Cell fate determination is critical for tissue development and cancer progression. Here, we demonstrated p63 is a critical regulator of the switch between squamous and neuroendocrine lineages in the esophagus. *p63* deletion promotes the differentiation of esophageal progenitor cells into neuroendocrine cells in both mouse and human esophageal progenitor cells. Conversely, ectopic ΔNp63α expression promotes the squamous differentiation of eNEC cells by directly binding to the squamous genes. Our study further revealed that p63 is epigenetically silenced by EZH2-mediated H3K27me3, which is accompanied by a reduction in H3K27ac levels. Treatment with EZH2 inhibitors leads to p63 expression and promotes squamous differentiation of eNECs. These results suggest that the lineage gatekeeping function of p63 is hijacked by eNECs to maintain the neuroendocrine identity and represents a potential target for epigenetic therapies.

p63 is a master regulator of basal cells, and ablation of *p63* leads to failed stratification of the epithelium in the developing esophagus (*13*). We showed here that p63 not only regulates the generation of the stratified epithelium, but also actively blocks non-squamous cell fate. These functionalities are conserved in human esophageal progenitor cells. The *p63* gene employs two different promoters to generate two major isoforms (*36*). Our findings indicate that ΔNp63α plays a more prominent role than TAp63α in preserving squamous cell fate. In line with this, ΔNp63α overexpression promotes squamous differentiation of eNEC cells. Conversely, ΔNp63α knockdown blocks the squamous differentiation of eNECs induced by EZH2 inhibition. p63^+^ basal cells are also abundantly present in the prostate where transdifferentiation towards neuroendocrine carcinoma occurs (*39*). Whether p63 plays a broader role in governing the shuttle between squamous and neuroendocrine cell fates in other tissues remains to be addressed.

The cell of origin for eNEC is not clear. Neuroendocrine cells are considered as the major cell of origin for small cell lung cancers (SCLC) (*40, 41*). Merkel cells are found in the middle portion of the human esophagus (*22*), and a clinical study revealed that eNEC arises primarily in the middle or lower third of the esophagus (*42*). However, the presence of Merkel cells in the esophagus of other species including rodents remains unknown. We previously reported that during esophageal-tracheal separation a subpopulation of epithelial progenitor cells originating from the trachea and lung translocate and incorporate into the esophagus as p63^+^ basal cells (*43*). It will be interesting to determine whether these cells of respiratory origin contribute to the formation of eNEC. Notably, the mutational and copy number variation signatures of eNEC are more similar to ESCC than SCLC (*44*), suggesting a possible common origin of eNEC and ESCC. In addition, eNEC was reported to arise from ESCC following combined treatment with chemotherapy and radiotherapy (*9*). Our study favors basal/squamous cells or their progenitors as one of the cells-of-origin for eNEC. Our findings further suggest that silencing of p63 likely represents a critical step in the squamous-to-neuroendocrine conversion during tumor progression. Such lineage transition seems reversible. Ectopic ΔNp63α expression was sufficient to bind and activate squamous genes in eNEC cells. Thus, epigenetic regulation of p63 expression could be key to the lineage plasticity that underlies esophageal intra-tumoral heterogeneity and enables cancer cells to adapt to niche- and drug-induced stressful environment.

eNEC is rare yet extremely aggressive, with high metastatic potential and poor prognosis (*24*). No standard therapy has been established for eNEC. Chemo- and radiation therapies approved for ESCC have proven to be largely ineffective against eNEC. Furthermore, recent multiomics profiling reveals that unlike ESCC, eNEC is poorly immune infiltrated, suggesting that it is a poor candidate for immune checkpoint inhibitors (*27*). Our results suggest that several epigenetic inhibitors, including the FDA-approved EZH2 and HDAC inhibitors, robustly activate squamous gene expression program in eNEC via the de-repression of p63. We observed a modest yet significant effect of EZH2 inhibition on the proliferation of eNEC organoids. Future studies are needed to assess the impact of the EZH2 inhibitors and other epigenetic therapies on the squamous trans-differentiation and growth of eNEC tumors *in vivo*. Importantly, we propose that the switch to squamous cell state by EZH2 inhibitors may sensitize eNEC to chemo-, targeted and immuno-therapies approved for ESCC, and these combination treatment strategies warrant further investigations in preclinical and clinical studies.

In summary, we identified an important role of p63 in repressing the differentiation of esophageal progenitor cells into neuroendocrine cells in mouse models and ESCs. Ectopic p63 expression is sufficient to promote the transdifferentiation of eNECs towards squamous cells. We further showed that p63 is silenced via EZH2-mediated epigenetic modification in eNEC cells. Accordingly, EZH2 inhibition is sufficient to activate p63 which orchestrates squamous gene expression in eNEC cells. Our findings therefore identify an important regulatory mechanism by the EZH2-p63 axis in determining squamous vs. neuroendocrine cell fates during normal esophageal development and malignant transformation.

## Materials and Methods

### Mice

*p63-CreERT2(*45*)* mice were maintained on a C57BL/6 and 129SvEv mixed background. We crossed *p63-CreERT2* mice to generate *p63 KO* embryos (*p63-CreERT2* homozygous). Mice were maintained in the animal facilities of Columbia University under a 12-hour light/12-hour dark cycle. All animal experiments were conducted according to the procedures approved by the Columbia University Institutional Animal Care and Use Committee.

### Clinical samples

Human esophageal small cell carcinoma samples were collected from The First Affiliated Hospital of Xi’an Jiaotong University, Xi’an, Shaanxi, China. This study was approved by the ethics commissions of the participating hospitals with written informed consent from the patients.

### hESC culture

*p63 KO* H9 cell lines were originally generated by Dr. Anthony Oro lab at Stanford University (*46*). H9 hESC were maintained on mitotically arrested MEF feeder cells in the medium composed of 80 ml of DMEM/F1, 20 ml of Knockout Serum Replacement, 1 ml of GlutaMAX, 1 ml of MEM non-essential amino acids solution, 0.7 µl of 2-mercaptoethanol, 0.2 ml of primocin and 20 ng/ml FGF2. Cells were maintained in an incubator of 5% CO2 at 37℃. Human hESC research was conducted under the approval of the Columbia University Human Embryonic and Human Embryonic Stem Research Committee.

### hESC differentiation into esophageal epithelial cells and organoids

We have reported a detailed protocol for esophageal differentiation of hESCs (*34, 35*). Serum free differentiation (SFD) medium was prepared as following: 150 ml of IMDM medium, 50 ml F12, 1.5 ml of 7.5% Bovine Albumin Fraction V Solution, 2 ml of GlutaMAX, 1 ml of N2, 2 ml of B27, 2 ml of Penicillin/Streptomycin and 50 µg/ml L-Ascorbic acid and 0.04 µl/ml 1-Thioglycerol. hESCs were differentiated into endodermal spheroids in SFD medium with 100 ng/ml Activin A, 0.5 ng/ml BMP2, 2.5 ng/ml FGF2 and 10 nM Y-27632 for 72 hours in 6-well low-attachment plates. Endoderm spheroids were disassociated into single cells and replated in fibronectin-coated 24-well plates and induced into foregut endoderm epithelial cells with 100 ng/ml Noggin and 10 mM SB431542 for 48 hours. To induce esophageal specification cells were continued to be cultured in 100 ng/ml Noggin and 10 mM SB431542 plus 100 ng/ml EGF for 4 days and 100 ng/ml EGF for another 14 days. Esophageal epithelial cells were then subjected to IF staining and qRT-PCR analysis. For esophageal organoid establishment, we embedded endodermal spheroids in Matrigel and cultured the organoids with SFD medium supplemented with 100 ng/ml Noggin, 10 mM SB431542 and 100 ng/ml EGF until day 53. Organoids were then harvested for immunostaining analysis.

### TYUC-1 cell culture and tumor organoid culture

TYUC-1 is the only available eNEC cell line that was originally established from a patient with esophageal small cell carcinoma (*47*). The cell line was maintained in DMEM/F12 (Corning) plus 10% serum and 1% penicillin/streptomycin in suspension culture. To establish 3D tumor organoids, TYUC-1 cells were digested with 0.05% Trypsin-EDTA at 37℃ for 15 minutes and disassociated with P1000 pipettes into single cells. 20,000 cells in 100 µl of medium were then mixed with 100 µl of Matrigel (Corning) and added to the 24-well inserts. Matrigel-embedded cells were allowed to solidify in a cell culture incubator at 37℃ for 30 minutes before medium was added. The organoid culture medium contained DMEM/F12 supplemented with 10% serum and 1% penicillin/streptomycin. Medium was refreshed every other day. To inhibit EZH2 activity, TYUC-1 cells or organoids were treated with either 2 μM EPZ-6438 or 2 μM GSK126 with DMSO as vehicle control.

### EZH2 shRNA lentivirus generation

To knockdown EZH2, two shRNAs targeting EZH2 (shEZH2-a: CCGG<u>CCCAACATAGATGGACCAAAT</u>CTCGAG<u>ATTTGGTCCATCTATGTTGGG</u>TTTTT G and shEZH2-b: CCGG <u>CGGAAATCTTAAACCAAGAAT</u>CTCGAG<u>ATTCTTGGTTTAA GA TTTCCG</u> TTTTTG) were designed and cloned into pLKO.1 vector (Addgene plasmid #10879) according to method by Moffat, *et al* (*48*). The control shRNA sequence was CCTAAGGTTAAGTCGCCCTCGCTCGAGCGAGGGCGACTTAACCTTAGG. Plasmids were amplified in *E. coli DH*5α and isolated using Qiagen QIAprep Spin Miniprep Kit (Qiagen) according to the manufacturer’s instructions. Lentivirus was produced with the packaging plasmids psPAX2 (Addgene plasmid #12260) and pMD2.G (Addgene plasmid #12259) in 293T cells. TransIT (Mirus) was used to transfect the plasmids. TYUC-1 cells cultured in suspension were infected with lentivirus, and 1 μM polybrene was added to improve the infecting efficiency. Cells were collected 24 hours later and resuspended for organoid culture as mentioned above. To knock down human TP63, siRNA targeting human TP63 (ON-TARGETplus, catalog no. L-003330-00-0005) and control siRNA (ON-TARGETplus, catalog no. D-001810-10-05) were used. siRNAs were transfected with Lipofectamine™ RNAiMAX (Thermo Fisher Scientific).

### Overexpression of TAp63 and ΔNp63 in TYUC-1 cells

The coding sequence of TAp63 and ΔNp63 was amplified with high-fidelity Taq polymerase using primers TAp63-EX-F: gaataccggtgcgctgccaccATGAATTTTGAAACTTCACGGTGTGCC and TAp63-EX-R: cgggatccCTCCCCCTCCTCTTTGATGC, ΔNp63-EX-F: gaataccggtctagagctgccaccATGGGCTCCGGCTCCTTGTACCTGGAAAACAATGCC and TAp63-EX-R. PCR products were digested with AgeI and BamHI, and cloned into the Tet-on plasmid TLCV2 (Addgene plasmid #87360) digested with the same enzymes. Lentivirus was packaged with the plasmids psPAX2 and pMD2.G in 293T cells. TYUC-1 cells were infected with lentivirus containing TAp63 or ΔNp63, with original empty vector plasmid as control. Infected cells were maintained in the culture medium supplemented with 1 μg/ml puromycin.

### Immunostaining

Antigen retrieval on tissue slides were performed using antigen unmasking solution (Vector laboratories #H-3301-250). Cells were fixed with 4% paraformaldehyde in 1xPBS for 10 minutes. Tissues or cells were treated with blocking solution composed of 1xPBS with 3% donkey serum and 0.3% Triton X-100 for 30 minutes. Primary antibodies diluted in blocking solution were added to the top of tissue slides and cells and incubated at 4℃ overnight. The next day, tissues and cells were washed with 1xPBS for three times before fluorophore-tagged secondary antibodies were added and incubated at room temperature for two hours. Tissues and cells were washed with 1xPBS for three times. Tissue slides were mounted with DAPI-containing mounting medium (Southern Biotech, #0100-20). Antibodies are summarized in table S1. Images were taken by Lecia DMi8 (Leica Microsystems) or LSM 700 laser scanning confocal microscope (Carl Zeiss). Representative images were generated by individually scanned images stitched with Leica Application Suite X software (Leica Microsystems) or Zen software (Carl Zeiss).

### RNA sequencing

The epithelium was isolated from the esophagus of E12.5 *p63 KO* (*p63-CreERT2* homozygous) mutants and littermate controls, and RNA was purified with PicoPure RNA Isolation Kit (Thermo Fisher Scientific). We determined RNA concentration with 2100 Bio-analyzer (Agilent Technologies), and then used Illumina TruSeq RNA prep kit (Illumina) to establish libraries which were sequenced with Illumina HiSeq400. Samples were multiplexed in each lane, which yields targeted number of single-end/pair-end 100 bp reads for each sample, as a fraction of 180 million reads for the whole lane. RTA (Illumina) was used for base calling and bcl2fastq (version 1.8.4) for converting BCL to fastq format, coupled with adaptor trimming. We mapped the reads to a reference genome (Mouse: UCSC/mm9) using Tophat (version 2.1.0) with 4 mismatches (--read-mismatches = 4) and 10 maximum multiple hits (--max-multihits = 10). The expression levels of each gene were presented as Fragments Per Kilobase of transcript per Million (FPKM) in this study. For RNA-sequencing of TYUC-1 cells, total RNA was extracted in TRIzol (Invitrogen) and precipitated in ethanol (DECON Labs). For RNA-sequencing of doxycycline-treated cells for induction of Tap63 and ΔΝp63 expression, libraries were prepared using NEBNext Ultra kit (New England Biolabs #E7490, #E7770, #E7335, #E7500) and sequenced using a Nextseq500/550 sequencer. Paired-end reads were obtained and mapped to the human genome assembly hg38 using HISAT2 (v2.1.0). The mapped reads count of each gene was measured by featureCounts (v1.6.1). For RNA-sequencing of EPZ-6438-treated cells, RNA samples were submitted to Columbia University Genome Center for library preparation, sequencing and bioinformatic analysis. For both experiments, differential gene expression was calculated by the R package DESeq2 (v1.28.0) and gene set enrichment analysis (GSEA) was performed by using GSEA software (v4.1.0) (*49*). Visualization was done using the ggplot2 R package.

### Quantitative Real-Time Polymerase Chain Reaction (qRT-PCR)

We used TRIzol to lyse cells or tissues and purified the RNA with RNeasy Mini Kit (QIAGEN). Reverse transcription was performed using the SuperScript III First-Strand SuperMix (Invitrogen). cDNA abundance was measured by real-time PCR using the iQ SYBR Green and StepOnePlus Real-Time PCR System (Applied Biosystems). The transcript levels of each gene were normalized to β-actin or GAPDH. Primer sequences are summarized in table S2.

### CUT&Tag

CUT&Tag was performed as described previously (*50*). In brief, 2×10^5^ cells were washed once with 1 ml of wash buffer (20 mM HEPES pH 7.5, 150 mM NaCl, 0.5 mM Spermidine (Sigma-Aldrich), 1x Protease inhibitor cocktail (Roche). Concanavalin A-coated magnetic beads (Bangs Laboratories) were washed twice with binding buffer (20 mM HEPES pH 7.5, 10 mM KCl, 1 mM MnCl2, 1 mM CaCl2). 10 μl/sample of beads were added to cells in 400 μl of wash buffer and incubated at room temperature for 15 min. Beads-bound cells were resuspended in 100 μl of antibody buffer (20 mM HEPES pH 7.5, 150 mM NaCl, 0.5 mM Spermidine, 0.06% Digitonin (Sigma-Aldrich), 2 mM EDTA, 0.1% BSA, 1× Protease inhibitor cocktail and incubated with H3K27me3 antibody (Cell Signaling Technology #9733), H3K27ac antibody (Active Motif #39134), BRD4 antibody (Epicypher #13-2003) or normal rabbit IgG (Cell Signaling Technology #2729) at 4 ℃ overnight on nutator. After being washed once with Dig-wash buffer (20 mM HEPES pH 7.5, 150 mM NaCl, 0.5 mM Spermidine, 0.05% Digitonin, 1x Protease inhibitor cocktail), beads-bound cells were incubated with 1 μl Guinea pig anti-rabbit secondary antibody (Antibodies Online ABIN101961) and 2 μl Hyperactive pA-Tn5 Transposase adapter complex in 100 μl Dig-300 buffer (20 mM HEPES-NaOH, pH 7.5, 0.5 mM Spermidine, 1x Protease inhibitor cocktail, 300 mM NaCl, 0.01% Digitonin) at room temperature for 1 h. Cells were washed three times with Dig-300 buffer to remove unbound antibody and Tn5 and then resuspended in 300 μl of tagmentation buffer (10 mM MgCl2 in Dig-300 buffer) and incubated at 37 °C for 1 h. 10 μl of 0.5 M EDTA, 3 μl of 10% SDS and 5 μl of 10 mg ml−1 Proteinase K were added to each sample and incubated at 50 °C for 1 h to terminate tagmentation. DNA was purified using chloroform isoamyl alcohol (Sigma Aldrich) and eluted with 25 μl ddH2O. For library amplification, 21 μl of DNA was mixed with 2µL i5 unique index primer (10 µM), 2 µL i7 unique index primer (10 µM) and 25 µL NEBNext® High-Fidelity 2X PCR Master Mix (NEB) and subjected to the following PCR program: 72 ℃, 5 min; 98 ℃, 30 sec; 13 cycles of 98 ℃, 10 sec and 63 ℃, 10 sec; 72 ℃, 1 min and hold at 10 ℃. To purify the PCR products, 1.1x volumes of pre-warmed Ampure XP beads (Beckman Coulter) were added and incubated at room temperature for 10 min. Libraries were washed twice with 80% ethanol and eluted in 20 μl of 10 mM Tris-HCl, pH 8. Libraries were sequenced on an NextSeq 550 platform (Illumina, 75 cycles High Output Kit v2.0) and 75-bp paired-end reads were generated. For H3K27me3 and H3K27ac, 2 µl of SNAP-ChIP K-MetStat panel and K-AcylStat panel nucleosomes (EpiCypher) respectively, were added as spike-in control at the primary antibody incubation step. For H3K27ac and BRD4 CUT&Tag were initially lightly fixed with 0.1% paraformaldehyde for 5 minutes and neutralized by 125 mM Glycine to preserve target stability.

### CUT&Tag data analysis

CUT&Tag reads of TYUC-1 cell samples were mapped to the mouse human assembly hg38 using Bowtie2 (v2.3.5.1, parameters: --local --very-sensitive-local --no-unal --no-mixed --no-discordant --phred33 -I 10 -X 700). Potential PCR duplicates were removed by the function "MarkDuplicates" (parameter: REMOVE_DUPLICATES=true) of Picard (v2.24.2). Genomic enrichments of CUT&Tag signals were generated using deeptools (v3.3.2, parameters bamCoverage -- normalizeUsing CPM --binSize 25 --smoothLength 100 --scaleFactor 1). Peaks were called using MACS2 and annotated by the R package ChIPSeeker (v1.28.3). For H3K27me3 and H3K27ac the number of reads for each barcode was counted to determine the scaling factor. Tracks were visualized using IGV.

### Chromatin chemical probe screen

For the chromatin chemical probe screening, 5×10^5^ TYUC-1 cells were plated and treated with EPZ-6438 (2 μM), SAHA (1 μM), Romidepsin (1 nM), JQ1 (500 nM), Seclidemstat (500 nM), EPZ5676 (10 μM), UNC0638 (1 μM), A485 (250 nM), MG149 (10 μM), PRT4165(10 μM) or DMSO. Medium was refreshed every two days.

### Quantification and statistical analysis

Data was presented as the mean ±SD using GraphPad Software Prism. Statistical significance was determined by Student’s t test. At least three biological replicates were included. P values of 0.05 or less were considered to be statically significant.

## Acknowledgments

We thank all the lab members in Que’s and Lu’s lab for discussion of the research and proofreading of the manuscripts. We thank Dr. Carmen Birchmeier-Kohler at the Max Delbrück Center for Molecular Medicine in Germany for providing anti-Insm1 antibody, and Dr. Alea A. Mills at Cold Spring Harbor Laboratory for providing *p63^loxp/loxp^* mice.

## Author contributions

Y.Z., D.K., H.L., C.L. and J.Q. conceived the study, designed the experiments, performed the experiments, analyzed the data, and drafted the manuscript. M.L., Y.F., M.J., X.C., S.S., H.H., J.S. and F.S. assisted in data collection and study design. P.Y., W.E.R., A.Z., A.E.O., A.K.R. and T.C.W. critically revised the manuscript for important intellectual content. All authors read and approved the final manuscript.

---

**Fig. S1.**
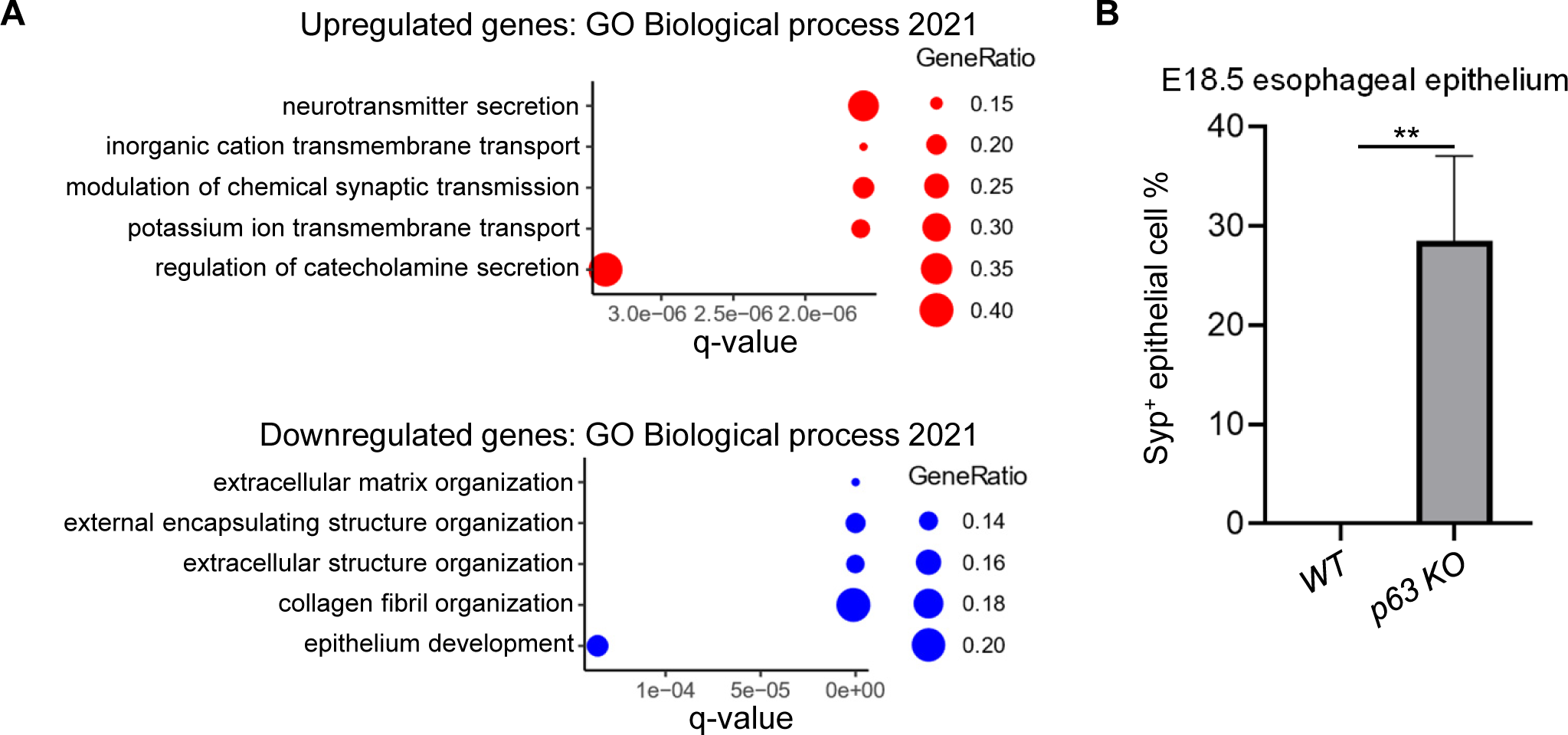
Gene ontology analysis of differentially expressed genes and quantification of Syp^+^ epithelial cells in the E18.5 esophagus. Related to Fig. 1. **(A)** Biological process gene ontology analysis of genes that are upregulated (red) and downregulated (blue) in the epithelium of E12.5 *p63 KO* esophagus. (B) Percentage of Syp^+^ cells of all epithelial cells in the E18.5 *p63 KO* and wildtype (WT) esophagus. **p < 0.01.

**Fig. S2.**
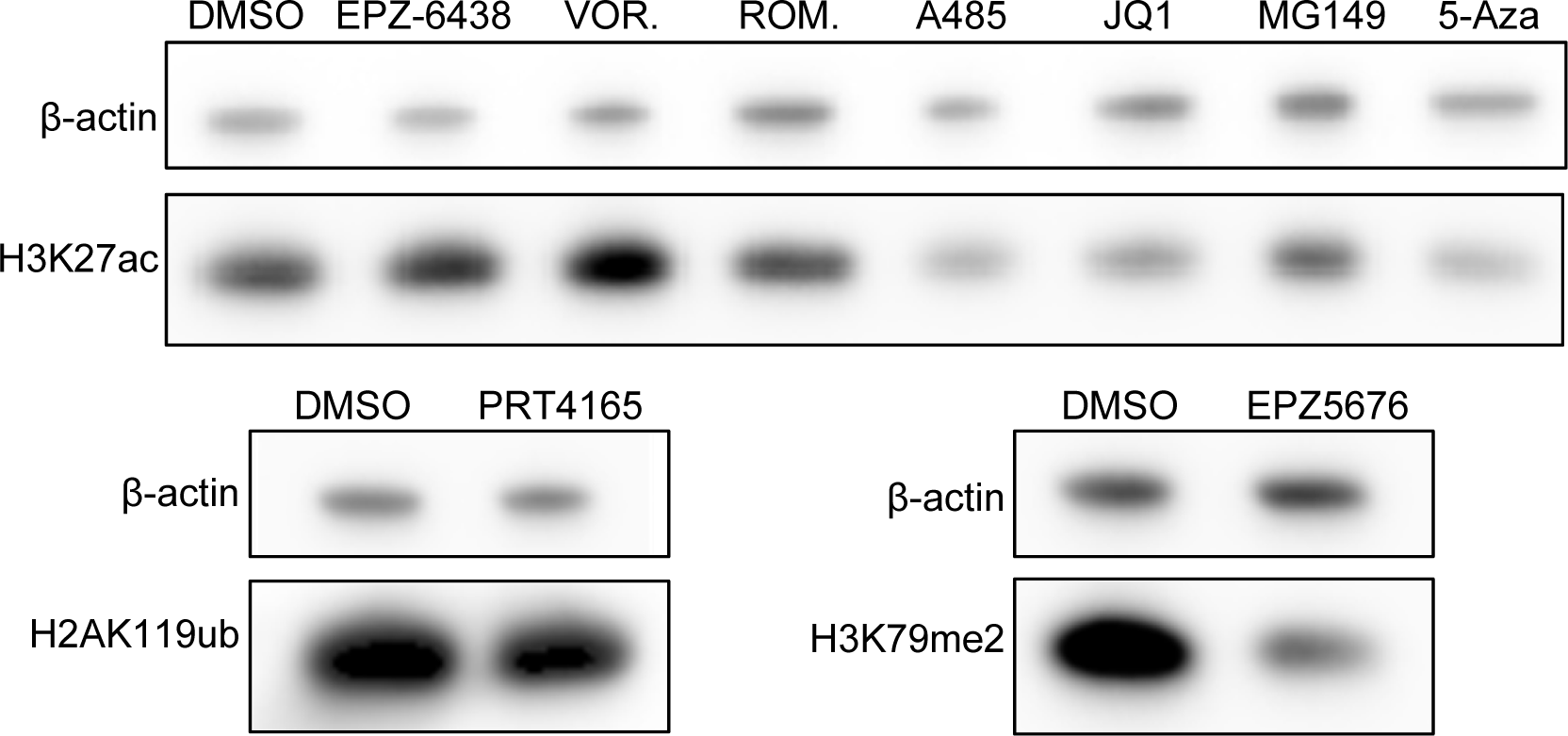
TYUC-1 cells treated with various inhibitors of epigenetic regulators. Related to Fig. 3. Western blots indicating levels of histone modifications upon treatment with the inhibitors of chromatin regulators. β-actin was included as loading controls. Vor, Vorinostat; ROM, Romidepsin; 5-AZA, 5-Azacytidine.

**Fig. S3.**
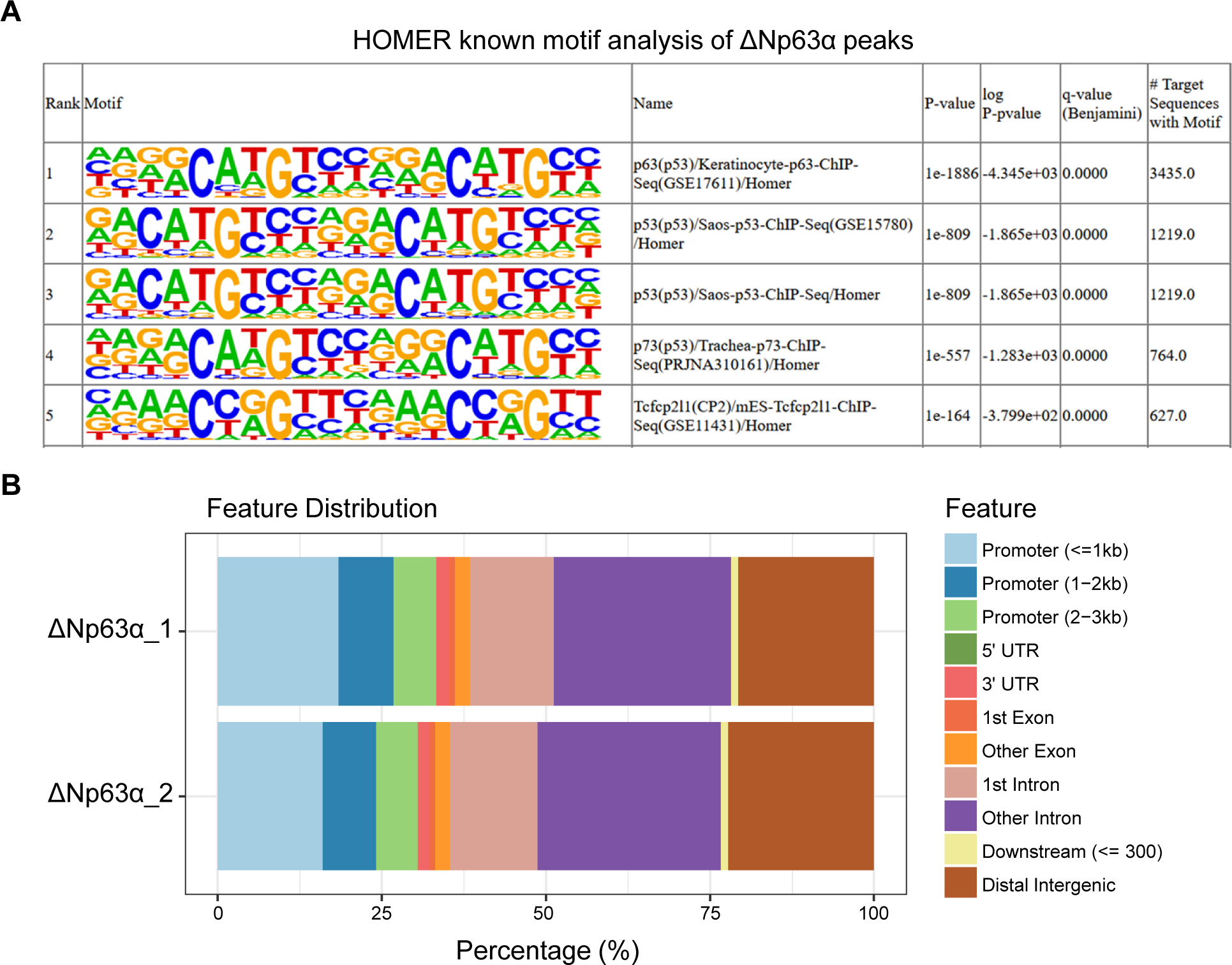
HOMER known motif analysis and genomic feature distribution of ΔNp63α biding sites in TYUC-1 cells overexpressing ΔNp63α. Related to Fig. 4. **(A-B)** HOMER known motif analysis of called peaks of ΔNp63α CUT&Tag upon induction of ΔNp63α (A); and distribution across genomic features (B; two replicates shown).

**Fig. S4.**
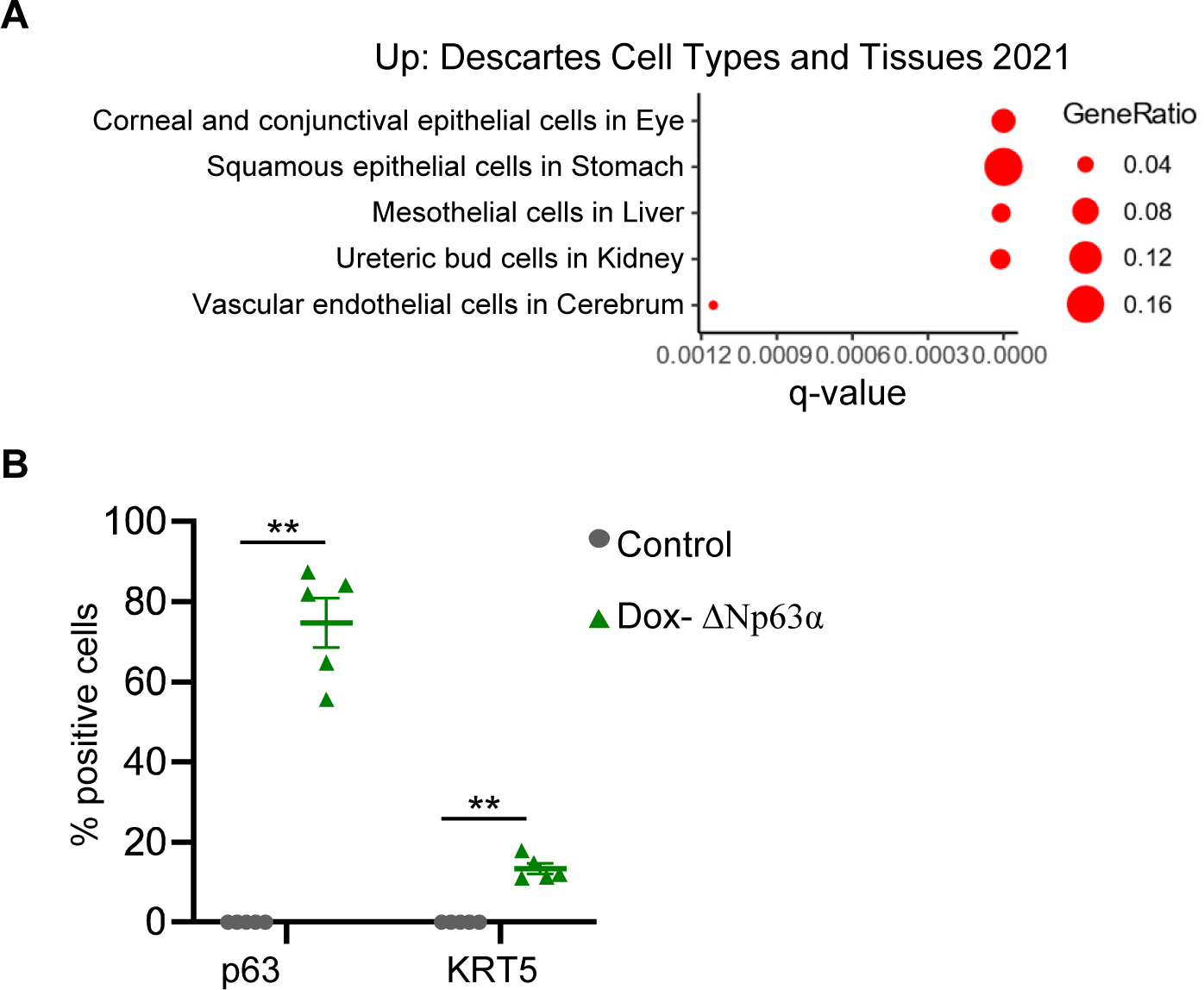
Gene ontology analysis of the upregulated genes and quantification of p63^+^ and KRT5^+^ cells in TYUC-1 cells overexpressing ΔNp63α. Related to Fig. 5. **(A)** “Cell Types And Tissues” gene ontology of upregulated genes in TYUC-1 cells upon induction of ΔNp63α expression by doxycycline for 6 days. **(B)** Quantification of p63**^+^** and KRT5**^+^** cells upon induction of ΔNp63α expression by doxycycline (Dox-ΔNp63α) for 6 days. **p < 0.01.

**Fig. S5.**
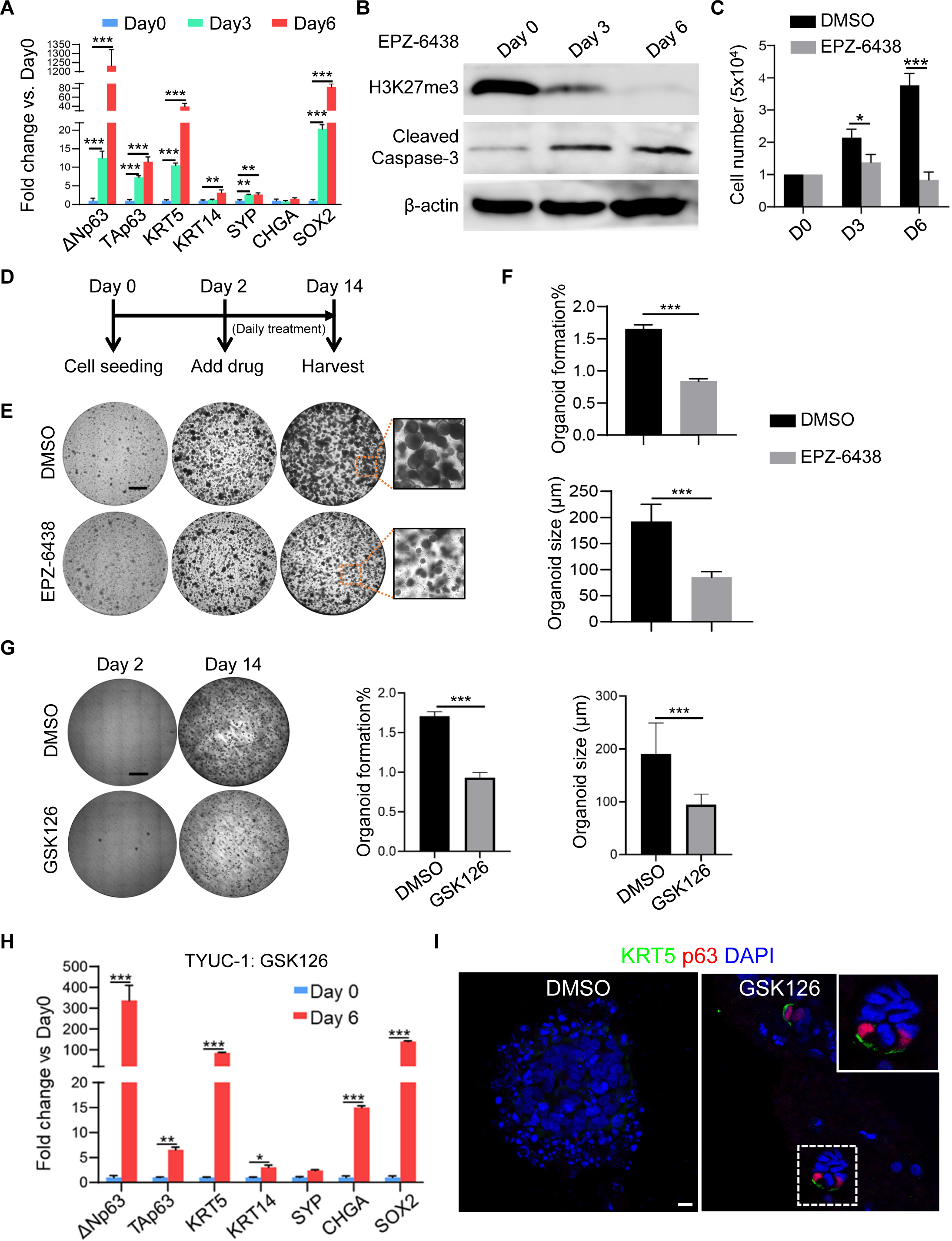
EZH2 inhibition promotes squamous differentiation of eNECs. Related to Fig. 6. **(A)** The transcript levels of the squamous cell markers ΔNp63, TAp63, KRT5, KRT4, SOX2, and the neuroendocrine cell markers SYP and CHGA in TYUC-1 cells treated with the EZH2 inhibitor EPZ-6438 for 3 or 6 days. **p < 0.01, ***p < 0.001. **(B)** Western blot indicating the protein levels of H3K27me3 and Cleaved Caspase-3 in TYUC-1 treated with EPZ-6438 for 3 or 6 days. The protein levels were determined by immunoblotting. **(C)** TYUC-1 cell proliferation was significantly reduced following treatment with EPZ-6438 for 3 or 6 days. *p < 0.05, ***p < 0.001. **(D)** Schematics showing 3D culture of TYUC-1 cells in Matrigel. Organoids were treated with either DMSO or 2 μM EPZ-6438 starting from day 2 and samples were collected at day 14. **(E)** Representative images of TYUC-1 organoids treated with either DMSO or EPZ-6438. **(F)** Formation efficiency and sizes of TYUC-1 organoids treated with DMSO or EPZ-6438. ***p < 0.001. **(G)** Representative images, formation efficiency and sizes of TYUC-1 organoids treated with DMSO or the EZH2 inhibitor GSK126. ***p < 0.001. **(H)** Transcript levels of squamous and neuroendocrine genes in TYUC-1 cells treated with the EZH2 inhibitor GSK126 for 6 days. *p < 0.05, **p < 0.01, ***p < 0.001. **(I)** Immunofluorescence staining images of TYUC-1 organoids treated with either DMSO or GSK126. Scale bars: 1mm in (E) & (G), 20 µm in (I).

**Fig. S6.**
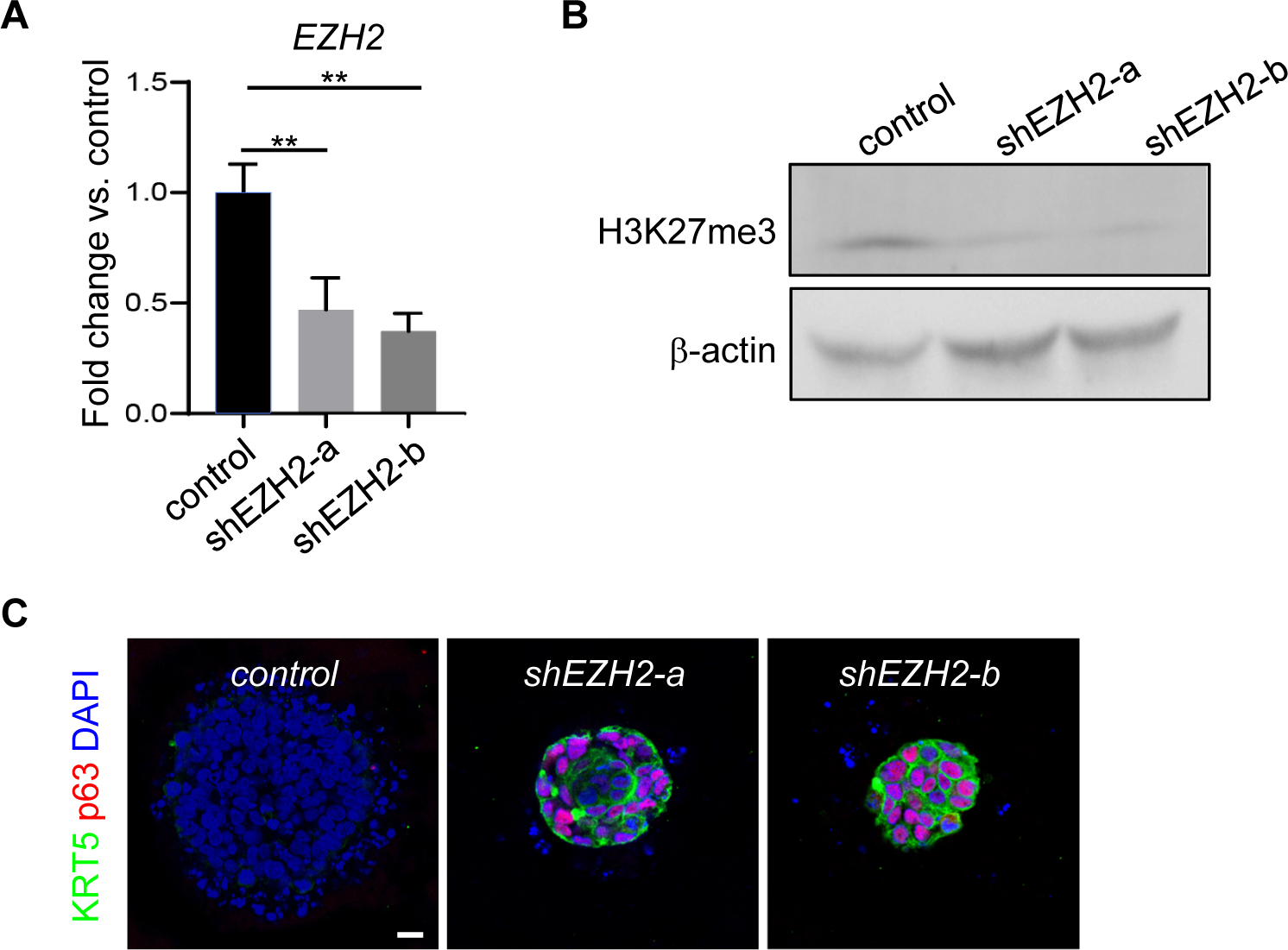
EZH2 knockdown promotes squamous differentiation of eNECs. Related to Fig. 6. **(A)** *EZH2* transcript levels were decreased in TYUC-1 cells infected with lentivirus containing two independent *EZH2* shRNAs. **p < 0.01. **(B)** Protein levels of H3K27me3 were decreased in TYUC-1 cells infected with lentivirus containing *EZH2* shRNAs. **(C)** Immunostaining of p63 and KRT5 in TYUC-1 organoids infected with lentivirus containing EZH2 shRNAs. Scale bar: 20 µm.

**Table S1.**
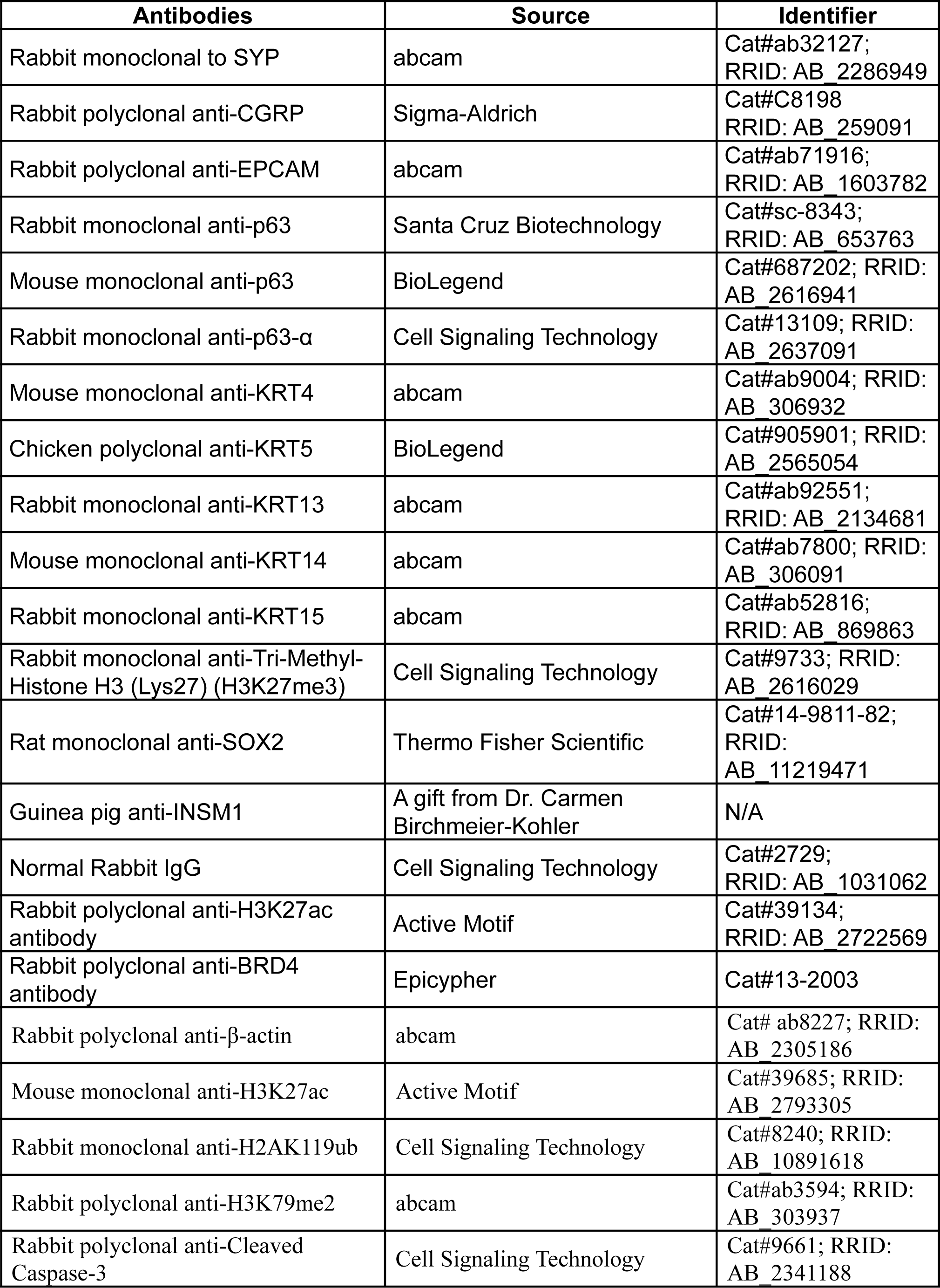
List of Antibodies.

**Table S2.**
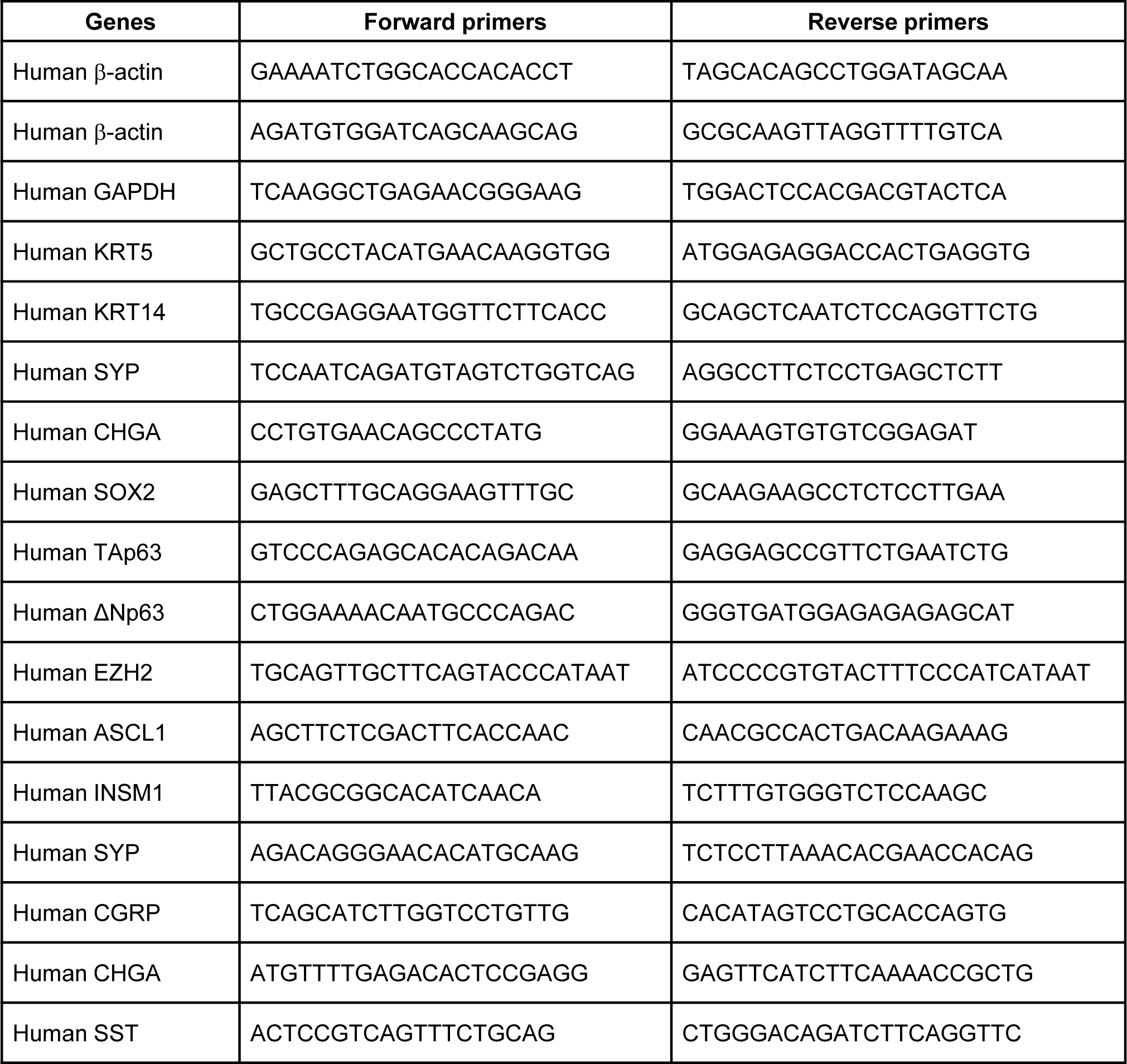
qRT-PCR primer sequences.

